# Production of the cobalamin lower ligand intermediate, α-ribazole, by a diatom under cobalamin-deplete conditions

**DOI:** 10.1101/2024.12.03.626653

**Authors:** Anna K. Gleason, Catherine C. Bannon, Erin M. Bertrand

**Author notes:** Max Planck Institute for Marine Microbiology, Bremen, Germany (Catherine C. Bannon). **Author Contribution Statement:** **AKG:** Conceptualization, Methodology, Investigation, Formal Analysis, Visualization, Data Curation, Writing – Original Draft, Writing – Review & Editing **CCB:** Conceptualization, Methodology, Supervision, Writing – Original Draft, Writing – Review & Editing. **EMB:** Conceptualization, Methodology, Supervision, Funding Acquisition, Resources, Writing – Original Draft, Writing – Review & Editing.

## Abstract

Cobalamin, or vitamin B_12_, is a micronutrient required by half of all surveyed phytoplankton but produced only by select bacteria and archaea. Cobalamin influences community composition and primary productivity in various regions of the ocean and has been shown to be a critical currency in microbial interactions. Many marine microorganisms can salvage cobinamide and remodel pseudocobalamin, to generate cobalamin using the lower ligand 5,6-dimethylbenzimidazole (DMB), or the ribosylated form, α-ribazole. The sources of the cobalamin lower ligand and intermediate remain poorly characterized but are currently attributed to prokaryotes alone. Here, we grew axenic cultures of a recently isolated *Thalassiosiraceae* diatom over a range of cobalamin concentrations (0, 0.75, 10, 100 pM). Then, using mass spectrometry, we quantified cobalamin and α-ribazole in both particulate (cellular) and dissolved (media) phases. We found that this diatom is a facultative cobalamin consumer: it takes up cobalamin when available but can survive in its absence. Strikingly, under low cobalamin availability, we demonstrate that the diatom produces α-ribazole. This highlights eukaryotic production as a source of the molecule and suggests that salvaging and remodeling by phytoplankton may be less dependent on prokaryotes than previously imagined. We detected α-ribazole in the spent media during stationary phase, highlighting eukaryotic production of the molecule as a public good. We provide evidence that α-ribazole is produced by this diatom as a mechanism to cope with low cobalamin availability, suggesting that even amongst organisms that don’t absolutely require it, cobalamin use has important consequences for fitness.

**Importance Section:** Cobalamin (vitamin B_12_) is a scarce micronutrient in the ocean that impacts the composition and activity of marine microbial communities. Microbes employ a range of strategies to cope with low cobalamin availability, including remodeling cobalamin intermediates and alternatives into cobalamin when the lower ligand is available. Here we show that a recently isolated *Thalassiosiraceae* diatom doesn’t need cobalamin to survive but will take it up and use it when available. Additionally, the diatom produces the lower ligand (α-ribazole) under low cobalamin conditions. The production of α-ribazole was previously thought to be restricted to bacteria and archaea, so these results suggest that eukaryotes could play a larger role in cobamide remodeling than previously imagined. This work also demonstrates that even organisms that don’t absolutely require cobalamin for survival have adopted metabolic strategies to cope with its absence, emphasizing the importance of this vitamin for phytoplankton metabolism.

## Main text

Cobalamin is a structurally complex micronutrient composed of a central cobalt-containing corrin ring, a lower (α) ligand in which 5,6-dimethylbenzimidazole (DMB) is incorporated as α-ribazole within the nucleotide loop, and an upper (β) ligand of either Ado-, Me-, OH-, or CN-. The biologically active forms of cobalamin are Ado- and Me-B_12_ (1) while CN-B_12_ is largely synthetic, and OH-B_12_ is the more stable, enzymatically inactive form (2–4). The biosynthesis of cobalamin requires over 30 enzymatic steps and is only completed by select bacteria and archaea (5,6) yet cobalamin is absolutely required by half of all surveyed eukaryotic phytoplankton and many heterotrophic bacteria, making them cobalamin auxotrophs (7,8). Eukaryotic phytoplankton that are not auxotrophs use the vitamin if it’s available, despite the fact that they can survive without it (9). However, the influence of cobalamin on these non-auxotroph facultative consumers is rarely considered.

Phytoplankton and many bacteria are capable of producing cobalamin from cobalamin-related compounds, including cobinamide and pseudocobalamin, using the lower ligand (6,10–13). However, a key remaining knowledge gap is the source of the lower ligand within in the ocean, which is currently assumed to be synthesis by prokaryotes and abiotic degradation of cobalamin, but has not been thoroughly investigated (4,10,14). This study leverages targeted mass spectrometry to confirm the cobalamin non-auxotrophic status of the diatom *Thalassiosiraceae* GTAN1 and document its production of α-ribazole under cobalamin deprivation.

We grew batch cultures of *Thalassiosiraceae* GTAN1, a diatom recently isolated from the Northwest Atlantic Ocean (supplemental methods), in axenic biological triplicates under a range of cobalamin concentrations (0, 0.75, 10, 100 pM CN-B_12_) (Fig 1A, B). No bacterial contamination was observed via plating techniques and flow cytometry (supplemental methods, Fig S1). After about 60 generations of growth in the respective treatments, we calculated growth rate, which did not significantly differ across these treatments (Fig 1B, p > 0.05) confirming that the diatom is not a cobalamin auxotroph.

**Figure 1:**
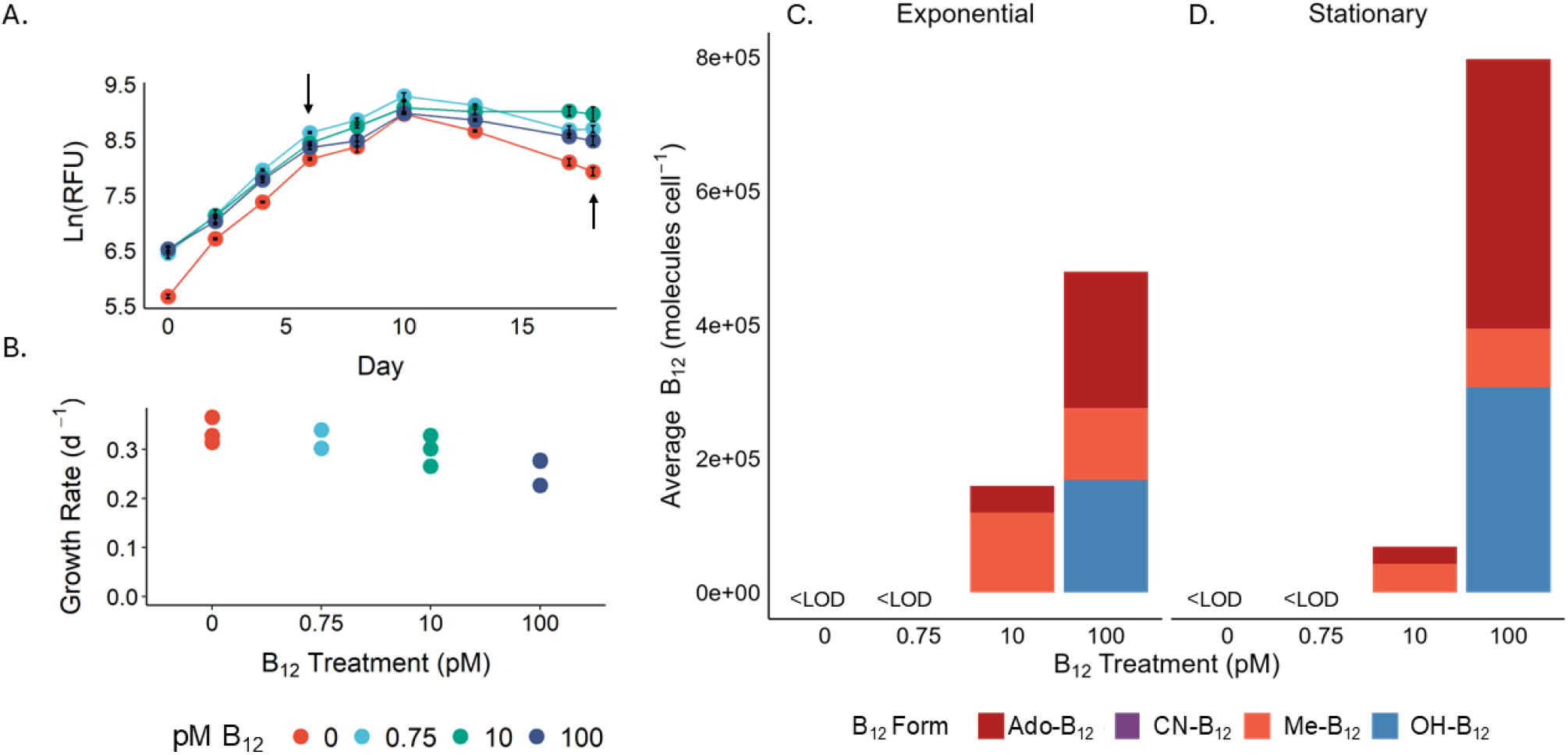
*Thalassiosiraceae* GTAN1 growth rates are not influenced by cobalamin availability, but the diatom will take up cobalamin when its available. (A) Growth curves (ln [RFU]) in different B_12_ treatments, arrows represent harvest dates for mass spectrometry analysis (B) growth rates (d^-1^) under different CN-B_12_ concentrations (0, 0.75, 10, 100 pM). Average particulate cobalamin content in biological replicates (n=3) in the diatom, with each B_12_ form (molecules cell^-1^) shown, measured during (C) exponential and (D) stationary phase. <LOD indicates values below the limit of detection.

We then quantified particulate cobalamin (>0.22 um) in exponential and stationary phase (arrows, Fig 1A) through targeted mass spectrometry (supplemental methods). All cobalamin forms were below the limit of detections (LOD) (Table S3) in the 0 and 0.75 pM treatments in the particulate samples in both exponential and stationary phase (Fig 1C, D). We observed that the CN-B_12_ taken up was converted into the biologically active forms, Me-B_12_ and Ado-B_12_ in cultures grown in 10 pM CN-B_12_. OH-B_12_, the enzymatically inactive intermediate, was found in cells grown in 100 pM CN-B_12_ treatment (Fig 1C, D), suggesting that this may be the dominant form that accumulates in cells during times of excess. Total cobalamin quotas ranged from 1.3 to 7.6 x 10^5^ molecules cell^-1^ across the 10 and 100 pM treatments (Fig 1C,D). These quotas are comparable to the other available diatom cobalamin quotas, reported for *T. pseudonana*, a cobalamin auxotroph. When grown in 1 pM and 200 pM CN-B_12_, *T. pseudonana* had total cobalamin quotas of 1.23 x 10^5^ and 3.4 x 10^5^ molecules per cell respectively (11). This result suggests that this diatom is a facultative consumer of cobalamin which will take up and use cobalamin when available and emphasizes that facultative consumers can be important sinks of cobalamin in the ocean.

During the targeted quantification of cobalamin in particulate samples, we also monitored various b-vitamins and vitamers (Table S1) including the cobalamin lower ligand of cobalamin. To our surprise, α-ribazole was measured above the limit of detection in all treatments in stationary phase and in select 0 and 0.75 pM samples in exponential phase (Fig 2A, B, Table S4). A-ribazole was not added to the media, and cobalamin was not detected in the 0 and 0.75 pM treatments (Fig 1C, D), meaning it was not a degradation product of supplied CN-B_12_. Production of α-ribazole was significantly higher (p-value < 0.05, Table S7) in 0 and 0.75 pM B_12_ treatments than in 10 pM or 100 pM B_12_ treatments in stationary phase (Fig 2B). Cellular quotas of α-ribazole ranged from below the limit of detection to 5.36 x 10^6^ molecules per cell (Figure 2A, B, Table S3). To our knowledge, there are no cellular quotas of lower ligand in literature to date. In the dissolved phase, α-ribazole was measured and quantified during stationary growth for the 0 pM treatment at concentrations (pM) approximately 40 times greater than in particulate samples, samples were not collected for other treatments and time points (Fig 2C).

**Figure 2:**
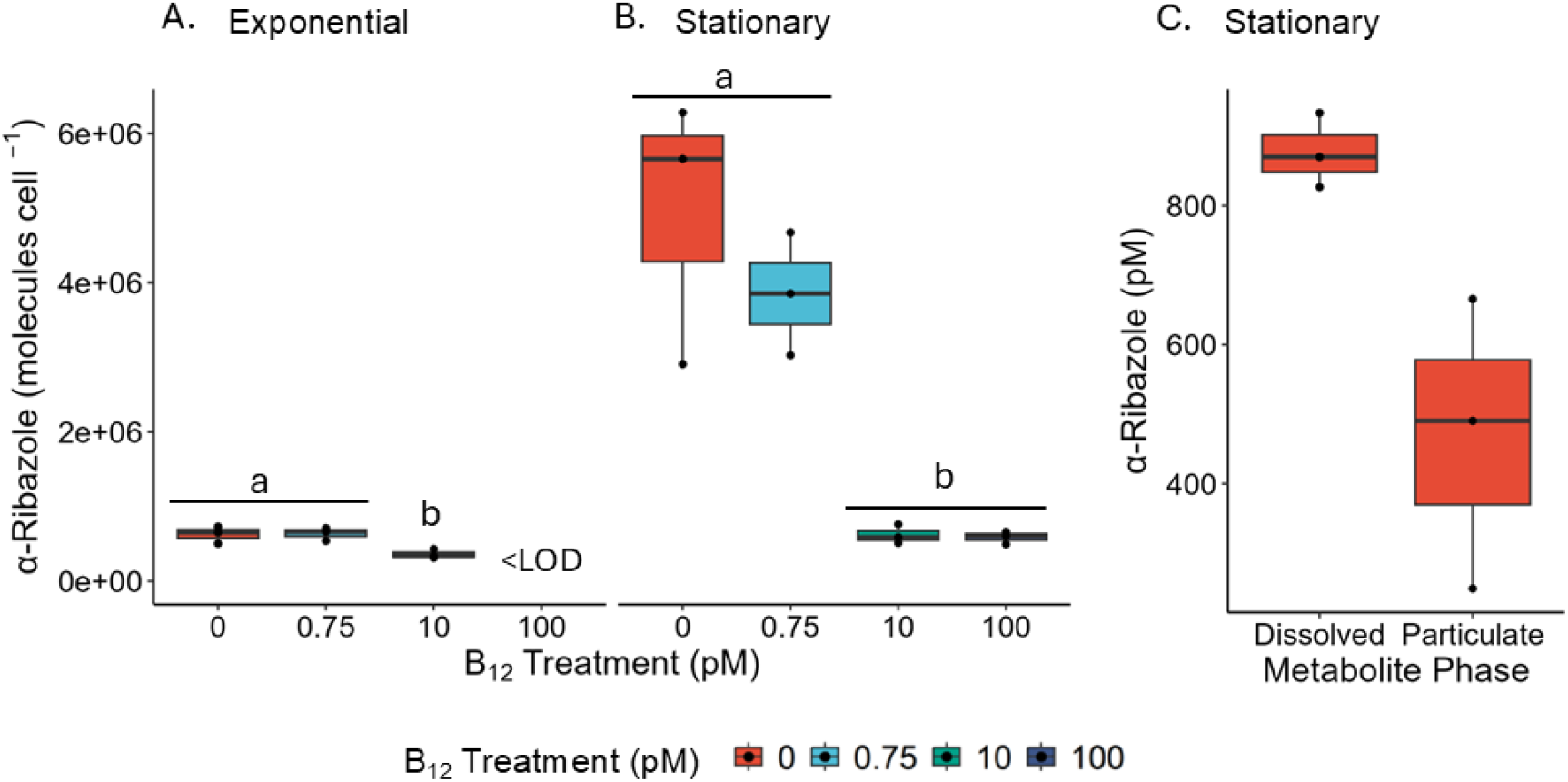
*Thalassiosiraceae* GTAN1 produces α-ribazole, the lower ligand of B_12_, which is detected in the media in low B_12_ treatments. Particulate α-ribazole quotas (molecules cell^-1^) in (A) exponential and (B) stationary phase. Different letters indicate statistically significant differences (p-value < 0.05) between means of biological replicates based on the post-hoc Tukey’s Test. (C) Concentrations of dissolved and particulate α-ribazole (pM) in stationary phase in 0 pM B_12_ treatment. Points represent biological replicates (n =3), points below limit of detection (<LOD) are not shown.

To date, two possible DMB biosynthesis pathways have been identified, the first being the BluB enzyme (5,6-dimethylbenzimidazole synthase, EC:1.13.11.79) which generates DMB from vitamin B_2_ (riboflavin) (15). Following its production by BluB, DMB is ribosylated into α-ribazole via the CobT pathway (16,17). An investigation into the horizontal gene transfer of cobalamin related genes in diatoms revealed that of the nine examined diatoms BluB was encoded in the genomes of *Fragiliariopsis cylindrus, Pseudo-nitzschia multistriata*, and *Pseudo-nitzschia multiseries* (18). The second pathway was identified in the obligate anaerobe, *Eubacterium limosum* where the bzaABCDE operon facilitates DMB production (19). However, mounting evidence suggests that there may be unannotated DMB biosynthesis pathways. For example, cultivation of *Rugeria pomeryoi* detected α-ribazole despite the bacterium lacking annotated genomic pathways for production of the molecule (20). While this measured α-ribazole may have originated from degradation of cobalamin provided in the media, the molecule was measured to be 3x greater in response to DMSP addition, suggesting that this bacterium may possess an unannotated biosynthesis pathway. DMB is not known to perform additional cellular functions other than acting as the lower ligand of cobalamin (11). While we are unable to present genomic evidence for lower ligand synthesis in*Thalassiosiraceae* GTAN1 currently, we offer the first measurements of production of the molecule and provide functional evidence for eukaryotic lower ligand synthesis.

*Thalassiosiraceae* GTAN1 may be producing α-ribazole under low cobalamin availability in order to generate cobalamin through subsequent salvaging and remodeling, consistent with the notion that even facultative consumer phytoplankton employ molecular mechanisms to cope with cobalamin deprivation (9). Pseudocobalamin remodeling has been demonstrated in non-auxotrophs and auxotrophs alike, but previously only when eukaryotes were supplied with exogenous DMB (10). This discovery suggests that some eukaryotes may not be as dependent on exogenous sources of the lower ligand for remodeling as once thought. The generation of dissolved lower ligand suggests that this diatom may contribute to cobalamin availability in the ecosystem both by conducting remodeling itself, and by producing α-ribazole which other community members can use as a common good to remodel.

There is increasing evidence for the potential of cobamide remodeling and cobinamide salvaging to significantly contribute to cobalamin availability in the ocean. Along the Scotian Shelf in the Northwest Atlantic Ocean, metagenomic and targeted metabolite measurement work points toward increased remodeling activity in the fall compared to the spring (21,22). The importance of remodeling is not limited to this region: the addition of α-ribazole was found to influence prokaryotic and protist communities in multiple regions of the Pacific Ocean (23). Recently, ligand cross-feeding including the exchange of α-ribazole, was observed to not only facilitate complex microbial interactions but to increase cobalamin availability, and is hypothesized to contribute significant amounts of cobalamin to ecosystems (13). This mounting evidence points towards the importance of cobamide remodeling for cobalamin availability, and the crucial role that DMB plays in this process. The identification of DMB production by this diatom provides evidence that we need to consider eukaryotic DMB production in our investigations into the role of remodeling in ecosystems.

## Acknowledgements

This work was made possible by NSERC Discovery Grant RGPIN2015-05009, Simons Foundation Grant 504183, Simons Foundation CBIOMES Award ID 1001702, Canada Research Chair Support to EMB, NSERC CGS-D to CB and a Nancy Witherspoon Memorial Summer Research Award awarded to AKG. The authors are grateful to the Bedford Institute of Oceanography, Fisheries and Oceans Canada Atlantic Zone Monitoring Program (AZMP) for access to the sea which allowed for the isolation of the diatom used in this study.

## Data Availability Statement

The datasets are available in the supplemental information.

## References

1. Banerjee R, Ragsdale SW. The Many Faces of Vitamin B 12 : Catalysis by Cobalamin-Dependent Enzymes. Annu Rev Biochem. 2003 Jun;72(1):209–47. doi:10.1146/annurev.biochem.72.121801.161828

2. Roth J, Lawrence J, Bobik T. COBALAMIN (COENZYME B12 ): Synthesis and Biological Significance. Annu Rev Microbiol. 1996 Oct;50(1):137–81. doi:10.1146/annurev.micro.50.1.137

3. Warren MJ, Raux E, Schubert HL, Escalante-Semerena JC. The biosynthesis of adenosylcobalamin (vitamin B12). Nat Prod Rep. 2002 Jul 18;19(4):390–412. doi:10.1039/b108967f

4. Bannon CC, Mudge EM, Bertrand EM. Shedding light on cobalamin photodegradation in the ocean.

5. Rodionov DA, Vitreschak AG, Mironov AA, Gelfand MS. Comparative Genomics of the Vitamin B12 Metabolism and Regulation in Prokaryotes. Journal of Biological Chemistry. 2003 Oct;278(42):41148–59. doi:10.1074/jbc.M305837200

6. Shelton AN, Seth EC, Mok KC, Han AW, Jackson SN, Haft DR, et al. Uneven distribution of cobamide biosynthesis and dependence in bacteria predicted by comparative genomics. ISME J. 2019 Mar;13(3):789–804. doi:10.1038/s41396-018-0304-9

7. Croft MT, Lawrence AD, Raux-Deery E, Warren MJ, Smith AG. Algae acquire vitamin B12 through a symbiotic relationship with bacteria. Nature. 2005 Nov;438(7064):90–3. doi:10.1038/nature04056

8. Sañudo-Wilhelmy SA, Cutter LS, Durazo R, Smail EA, Gómez-Consarnau L, Webb EA, et al. Multiple B-vitamin depletion in large areas of the coastal ocean. Proceedings of the National Academy of Sciences. 2012 Aug 28;109(35):14041–5. doi:10.1073/pnas.1208755109

9. Bertrand EM, Allen AE, Dupont CL, Norden-Krichmar TM, Bai J, Valas RE, et al. Influence of cobalamin scarcity on diatom molecular physiology and identification of a cobalamin acquisition protein. Proc Natl Acad Sci USA. 2012 Jun 26;109(26). doi:10.1073/pnas.1201731109

10. Helliwell KE, Lawrence AD, Holzer A, Kudahl UJ, Sasso S, Kräutler B, et al. Cyanobacteria and Eukaryotic Algae Use Different Chemical Variants of Vitamin B12. Current Biology. 2016 Apr;26(8):999–1008. doi:10.1016/j.cub.2016.02.041

11. Heal KR, Qin W, Ribalet F, Bertagnolli AD, Coyote-Maestas W, Hmelo LR, et al. Two distinct pools of B 12 analogs reveal community interdependencies in the ocean. Proc Natl Acad Sci USA. 2017 Jan 10;114(2):364–9. doi:10.1073/pnas.1608462114

12. Ma AT, Tyrell B, Beld J. Specificity of cobamide remodeling, uptake and utilization in Vibrio cholerae. Molecular Microbiology. 2020 Jan;113(1):89–102. doi:10.1111/mmi.14402

13. Wienhausen G, Moraru C, Bruns S, Tran DQ, Sultana S, Wilkes H, et al. Ligand cross-feeding resolves bacterial vitamin B12 auxotrophies. Nature. 2024 May 23;629(8013):886–92. doi:10.1038/s41586-024-07396-y

14. Helliwell KE. The roles of B vitamins in phytoplankton nutrition: new perspectives and prospects. New Phytologist. 2017 Oct;216(1):62–8. doi:10.1111/nph.14669

15. Taga ME, Larsen NA, Howard-Jones AR, Walsh CT, Walker GC. BluB cannibalizes flavin to form the lower ligand of vitamin B12. Nature. 2007 Mar;446(7134):449–53. doi:10.1038/nature05611

16. Anderson PJ, Lango J, Carkeet C, Britten A, Kräutler B, Hammock BD, et al. One Pathway Can Incorporate either Adenine or Dimethylbenzimidazole as an α-Axial Ligand of B12 Cofactors in Salmonella enterica. J Bacteriol. 2008 Feb 15;190(4):1160–71. doi:10.1128/JB.01386-07

17. Crofts TS, Hazra AB, Tran JL, Sokolovskaya OM, Osadchiy V, Ad O, et al. Regiospecific Formation of Cobamide Isomers Is Directed by CobT. Biochemistry. 2014 Dec 16;53(49):7805–15. doi:10.1021/bi501147d

18. Vancaester E, Depuydt T, Osuna-Cruz CM, Vandepoele K. Comprehensive and Functional Analysis of Horizontal Gene Transfer Events in Diatoms. Battistuzzi FU, editor. Molecular Biology and Evolution. 2020 Nov 1;37(11):3243–57. doi:10.1093/molbev/msaa182

19. Hazra AB, Han AW, Mehta AP, Mok KC, Osadchiy V, Begley TP, et al. Anaerobic biosynthesis of the lower ligand of vitamin B _12_. Proc Natl Acad Sci USA. 2015 Aug 25;112(34):10792–7. doi:10.1073/pnas.1509132112

20. Johnson WM, Kido Soule MC, Kujawinski EB. Evidence for quorum sensing and differential metabolite production by a marine bacterium in response to DMSP. ISME J. 2016 Sep;10(9):2304–16. doi:10.1038/ismej.2016.6

21. Bannon C, White PL, Rowland E, More KJ, Gleason A, Roberts M, et al. Seasonal patterns in b-vitamins and cobalamin co-limitation in the Northwest Atlantic [Internet]. Microbiology; 2024 [cited 2024 Nov 20]. Available from: http://biorxiv.org/lookup/doi/10.1101/2024.11.10.622835 doi:10.1101/2024.11.10.622835

22. Soto MA, Desai D, Bannon C, LaRoche J, Bertrand EM. Cobalamin producers and prokaryotic consumers in the Northwest Atlantic. Environmental Microbiology. 2023 Mar 14;1462-2920.16363. doi:10.1111/1462-2920.16363

23. Wienhausen G, Dlugosch L, Jarling R, Wilkes H, Giebel HA, Simon M. Availability of vitamin B12 and its lower ligand intermediate α-ribazole impact prokaryotic and protist communities in oceanic systems. The ISME Journal. 2022 Aug 1;16(8):2002–14. doi:10.1038/s41396-022-01250-7

