## Supplemental Information for "Production of the cobalamin lower ligand intermediate, α-ribazole, by a diatom under cobalamin-deplete conditions"

### Supporting Information

**Document Includes:**

***Supporting methods***

Diatom isolation and identification
*Thalassiosiraceae GTAN1* culture acclimation
Experimental design
Bacterial contamination testing
Mass spectrometry
Growth rates and statistical analyses

**Tables S1-S7**

Table S1 Transition list for targeted analytes
Table S2 Slope and $R^2$ of calibration curves for targeted analytes in particulate phase
Table S3 Observed intracellular cobalamin and DMB in *Thalassiosiraceae GTAN1* (molecules x $10^3$ cell$^{-1}$)
Table S4 Observed intracellular cobalamin and DMB with standard error of the mean in *Thalassiosiraceae GTAN1* (attomole cell$^{-1}$).
Table S5 One-way ANOVA, Shapiro-Wilk Test, and Levene Test for the effects of cobalamin availability on alpha-ribazole concentration.
Table S6 Post-hoc Tuckey's Test for the effects of cobalamin availability on alpha-ribazole concentration.

**Figure S1**

Figure S1 Assessment of bacterial contamination in diatom cultures

### *Supplemental Methods*

#### Diatom isolation and identification

*Thalassiosiraceae* GTAN1, a centric diatom roughly 30 µm in diameter, was isolated from the Northwest Atlantic Ocean (NWA) on the Atlantic Zone Monitoring Program SEG B01 (Newfoundland AZMP, JC190, Nov. 2nd – Dec. 3rd, 2019, 46°35'00.0 N, 52°56'00.0). Serial dilution isolation was performed to obtain single eukaryotic cells. To obtain axenic conditions, 10 mL cultures were inoculated with 100 µg per mL streptomycin and 50 µg per mL gentamycin. After antibiotics were added, the cultures were put back into the incubator but protected from light for 48 hours. Cultures were then inoculated (dilution 1:10) into fresh media and assessed visually for growth. Once biomass was accumulated, bacterial contamination checks were performed by spreading ~ 50 µL cultures on ½ Marine Broth 1.3% agar plates grown at room temperature (25 °C).

Monoculture of *Thalassiosiraceae* GTAN1 was identified via 18S rRNA gene sequencing. DNA was extracted using a Blood and Tissue DNAeasy Qiagen kit with few modifications. 10 mL of culture was pelleted and resuspended in sterile F/2 media (1). To break the cells, 0.1 mg of 100 µm and 400 µm beads were added and vortexed for 1 minute. After AW2 wash, cultures were centrifuged at max speed for a minute. Samples were eluted in 2 x 100 µL $H_2O$, and warmed to 37 °C. DNA quantification was performed on a nanodrop, a blank of F/2 media was run, and 1 uL of DNA extract was run, DNA concentration was assessed to be 27.7 ng/ uL with a cleanliness score of 2. PCR mix (11x) for 20 µL reaction was as followed; 22 µL each eukA and eukB primers, 110 µL ecoTaq mix and 60.5 µL ultrapure water. This is equivalent to 0.5 µL template, 2 µL each primer and 10 µL Taq + water. PCR settings were as such; 96 °C for 2 mins, then 30 mins, 58 °C for 1 min, 72 °C for 2 minutes and 30 seconds, 72 °C for 10 minutes (x 34 cycles) then the cycle was paused at 4 °C. PCR reaction was confirmed on 1% gel agarose with sodium borate buffer (made from 10X stock) and RedSAFE. PCR products were sent to Genome Quebec for 18s rRNA gene sequencing. The resulting nearly full length 18s rRNA sequence (Supplemental dataset) was compared via blastn to ncbi nr on 28 Nov 2024 and identified as 99% identical to both *Thalassiosira* minima strain CCMP 991 18S small subunit ribosomal RNA gene, partial sequence, Sequence ID: DQ093366.1; Identities: 1731/1733(99%) and *Bacterosira constricta* isolate SMDC01286 18S ribosomal RNA gene, partial sequence; Sequence ID: KT692951.1; Identities 1731/1733(99%). Both these species are members of the Thalassiosiraceae family, so we refer to our strain as *Thalassiosiraceae* GTAN1. GTAN is the culture naming convention in the Bertrand lab, referring to gta'n, the Mikmaw word for Ocean.

#### ***Culture*** **acclimation**

Cultures were grown at 10 °C in F/2 media adjusted to 0 pM, 0.75 pM and 10 pM of CN-$B_{12}$. Cultures were transferred (diluted 1:20) about every 14 days with fresh media for a total of 11 transfers (178 days). Growth curves were taken throughout this period by measuring relative fluorescence units (RFU) every third day. For this, 200 µL of culture were aliquoted onto a Corning Costar black 96-well plate and were read on a BioTek Synergy H1 microplate reader using an excitation wavelength of 440 and emission wavelength of 680 optimized to measure chlorophyll concentrations and reported as relative chlorophyll-a fluorescence units (RFU). Microscopy cell counts were compared to RFU to ensure that RFU was an accurate proxy for diatom cell counts and growth.

#### Experimental design

On the 11th culture transfer (day 160), *Thalassiosiraceae* GTAN1 was inoculated in triplicates into 30 mL of F/2 media at four different concentrations of cobalamin (CN-$B_{12}$): 0 pM, 0.75 pM, 10 pM, and 100 pM. 100 pM cultures were inoculated from 10 pM cultures, all other cultures were inoculated from the same cobalamin treatments. RFU was measured every other day as previously described. At mid exponential phase (day 6), 10 mL of each replicate cultures were harvested for particulate metabolite samples on nylon filters and stored at -80 °C until processing. During stationary phase (day 18), the remaining volume (about 12 mL) was harvested for particulate metabolite samples as previously described. In addition, dissolved samples were collected from cultures in 0 pM cobalamin conditions and stored in amber glass vials at -20 °C.

#### Bacterial contamination testing

Throughout the acclimation and growth, bacterial contamination of samples was tested via flow cytometry and microscopy. Briefly, 800 µL of each replicate was filtered using a 3 µm syringe filter and 198 µL of filtrate was aliquoted into a 96-well plate then stained with SYBR Green (1% v/v) and incubated in the dark for ten minutes. Bacterial cell counts of the filtrate were determined using a NovoCyte flow cytometer (Agilent Technologies, San Diego, CA), Parameters were set to a limit of 50 µL or 50,000 events. Before the exponential harvest, bacterial contamination checks were also performed by spreading ~ 35 µL cultures on ½ Marine Broth (Defico) 1.3% agar plates grown at room temperature (23 °C).

**Mass spectrometry**

*Particulate metabolite extraction*

Particulate metabolites were extracted as described in Heal et al (2), with minor modifications. Samples were extracted in a dark room using a red-light source to avoid photodegradation and kept on ice whenever possible. To lyse cells, 0.2 mL of each 100 µm and 400 µm silica beads were added to the bead beater tubes containing sample filters. 1 mL of ice-cold solvent mixture (40:20:20 acetonitrile: methanol: MQ water) was added and the mixture was agitated using a bead beater (MP Biomedicals) in 3 x 40 second pulses at 1800 rotations per minute (RPM) over a 20-minute period. The tubes were centrifuged briefly on a bench-top centrifuge and the supernatant was removed to a sterile tube. Centrifugation was repeated and supernatant was transferred. 300 µL of the extraction buffer was added to rinse the filter, samples were thoroughly vortexed then centrifuged and the supernatant was transferred to the 2 mL tube. Two more washes were completed with ice-cold methanol and combined with supernatant. Samples were placed in a vacufuge (Eppendorf Vacufuge plus) until dry and covered from the light. Once dry, samples were stored at -80 °C until analysis.

*Dissolved metabolite extraction*

Dissolved samples (0 pM $B_{12}$ treatment, stationary phase) were extracted on C18 solid phase extraction (SPE) HyperSep columns (Waters, 500 mg resin, 3 mL column volume). Columns were pre-conditioned with 2 mL of high-grade methanol and 2 mL of high-grade MQ water. Samples were loaded SPE columns using a vacuum manifold, maintaining a flow rate of 1 mL per minute. After samples were loaded, columns were washed of 3 x 1 mL MQ water then purged of water by running vacuum on high for 30 seconds. Samples were eluted using 1.7 mL high-grade methanol into sterile 2 mL centrifuge tubes then dried down in a vacufuge and stored in a -80 °C until analysis.

*Mass spectrometry analysis*

Both particulate and dissolved extracts were resuspended in 100 µL Buffer A (0.1 % formic acid and 2% acetonitrile) and briefly vortexed, then centrifuged at 4 °C at 15,000 rpm for 15 minutes to pellet cell debris, broken filter, and any beads that may have been transferred. 30 µL of clean extract was aliquoted to a clean HPLC vial and 10 µL of each sample were pooled into a clean 2 mL tube to create a quality control (QC) sample. All particulate samples were diluted 4-fold while dissolved samples were not diluted. Metabolites were quantified using a Dionex Ultimate-3000 LC system coupled to an electrospray ionization source of a TSQ Quantiva triple-stage quadrupole mass spectrometer (Thermo Science, Waltham, MA) using selected reaction monitoring (SRM) for a preselected list of transitions (Table S1) operating at the following settings: Q1 and Q3 resolution 0.7 (FWHM), 6 ms dwell time, CID Gas 2.5 mTorr, spray voltage in 3500 V positive ion mode, sheath gas 6, auxiliary gas 2, ion transfer tube temperature 325 °C, vaporizer temperature 100°C. Duplicate 5 µL injections were performed onto a nanoEase M/Z HSS T3, 300 µm x 150 mm, C18, 1.8 µm, 100 Å (PN186009249, Waters) column held at 50 °C and subject to a gradient of 4–99% B over 8 minutes. The total run time was 12 minutes.

Throughout the run, the QC mix was injected at the beginning and after every sample set in triplicates to monitor instrument response. Sample set specific calibration curves were performed in duplicate injections in 5, 10, and 50 fmol on analytical column for all forms of $B_{12}$ and DMB.

*Metabolite quantification*

Samples were analyzed using the software Skyline Daily (MacCoss Lab Software, University of Washington). Peaks were adjusted following lab standards and calibration curves. Dissolved DMB samples were adjusted for a percent recovery of 57% following Bannon et al, 2024 (3). Limit of detection (LOD) and limit of quantification (LOQ) were calculated as 3 and 5 times the standard deviation of the lowest concentration of calibration curves for each metabolite respectively and are reported in Table S2 and S3. Slopes of calibration curves were used to quantify the metabolite concentration in samples and $R^2$ values are reported in Table S2 and S3.

Cobalamin form samples with concentrations that fell below LOD were further visually inspected and analyzed in a batch-per-batch method based on the following criteria modified from previous studies (4,5). Concentrations were reported in samples that fell below the calculated LOD if, (i) the peak has the same retention time (± 0.2 min) as the authentic standard, (ii) two daughter fragments were present with co-occurring peaks, (iii) daughter fragments were present in the same order of intensity as authentic standard.

**Growth rates and statistical analyses**

Growth rate calculations and statistical analyses were performed in R/RStudio (Version 4.3.1). Specific growth rates ($d^{-1}$) were calculated using linear regression of ln RFU during the exponential growth phase using the R package segmented (6). Growth curves were created in R studio using the package ggplot2 (7). Statistical analyses were performed using R. Homogeneity of variance and normality of residuals assumptions were tested via Levene Test and Shapiro-Wilk Test respectively for each molecule in R (Table S6). If p-value > 0.05 assumptions were presumed to be met and a one-way ANOVA was used to determine significance followed by a Post-hoc Tuckey's Test to assess pairwise comparisons across cobalamin treatments (Table S7). If assumptions were not met for normality of residuals, a Kruskal Wallis test was completed to determine significance (Table S7). Independent t-tests were conducted to determine significant statistical differences between means for metabolite concentrations in particulate and dissolved phases. A significance level of $\alpha$ = 0.05 was used for each statistical test that was performed.

**Supplemental Tables**

**Table S1** Selected reaction monitoring mass spectrometry parameters for metabolites measured in this study.

| Analyte | Name | Precursor (m/z) | Product (m/z) | Collision Energy (eV) | Retention time (min) |
|---|---|---|---|---|---|
| Ado-$B_{12}$ | ado-cobalamin | 790.9 | 972.5 | 34.1 | 4.7 |
| | | 790.9 | 665.9 | 21.6 | 4.7 |
| Me-$B_{12}$ | methyl-cobalamin | 673.5 | 972.5 | 27.6 | 5.2 |
| | | 673.5 | 665.9 | 18.9 | 5.2 |
| OH-$B_{12}$ | hydroxy-cobalamin | 665.0 | 913.4 | 18.3 | 4.0 |
| | | 665.0 | 636.0 | 29 | 4.0 |
| CN-$B_{12}$ | cyano-cobalamin | 678.4 | 636.0 | 27 | 4.4 |
| | | 678.4 | 635.9 | 21 | 4.4 |
| α-ribazole | α-ribazole | 279.2 | 147.0 | 28 | 4.2 |
| | | 279.2 | 132.0 | 45 | 4.2 |
| | | 279.2 | 120.0 | 43 | 4.2 |

**Table S2** Slope and $R^2$ of calibration curves of targeted metabolites for particulate samples. LOD (fmol on HPLC column) calculated as 3 x (LOD) and 5 times (LOQ) the lowest concentration calibration curve. Reported in fmol on HPLC column and molecules $cell^{-1}$.

| Analyte | Slope | $R^2$ | LOD (fmol on HPLC column) | LOQ (fmol on HPLC column) | LOD (molecules $cell^{-1}$) | LOQ (molecules $cell^{-1}$) |
|---|---|---|---|---|---|---|
| Ado-$B_{12}$ | 4.13E+05 | 0.98 | 0.76 | 1.27 | 7.85E+04 | 1.31E+05 |
| Me-$B_{12}$ | 1.97E+04 | 0.97 | 0.84 | 1.41 | 8.70E+04 | 1.45E+05 |
| OH-$B_{12}$ | 3.78E+04 | 0.97 | 1.02 | 1.70 | 1.05E+05 | 1.75E+05 |
| CN-$B_{12}$ | 1.34E+04 | 0.99 | 0.77 | 1.28 | 7.90E+04 | 1.32E+05 |
| α-ribazole | 1.62E+05 | 0.97 | 1.57 | 2.62 | 1.62E+05 | 2.70E+05 |

**Table S3** Mean molecules per cell (± SE) of cobalamins and DMB in *Thalassiosiraceae GTAN1* (n=3).

| Growth Phase | Treatment | Ado-$B_{12}$ | Me-$B_{12}$ | OH-$B_{12}$ | CN-$B_{12}$ | α-ribazole |
|---|---|---|---|---|---|---|
| Exponential | 100 | 1.46E+05 ± 7.18E+04 | 1.07E+05 ± 2.85E+04 | 1.68E+05 ± 5.33E+04 | <LOD | <LOD |

| | | | | | | |
|---|---|---|---|---|---|---|
| Exponential | 10 | 4.01E+04 ± 1.18E+04 | 1.95E+05 ± 4.41E+04 | <LOD | <LOD | 3.60E+05 ± 2.91E+04 |
| Exponential | 0.75 | <LOD | <LOD | <LOD | <LOD | 6.37E+05 ± 3.80E+04 |
| Exponential | 0 | <LOD | <LOD | <LOD | <LOD | 6.28E+05 ± 4.97E+04 |
| Stationary | 100 | 3.82E+05 ± 4.52E+04 | 9.07E+04 ± 9.62E+03 | 2.85E+05 ± 5.57E+04 | <LOD | 5.87E+05 ± 3.72E+04 |
| Stationary | 10 | 2.58E+04* | 5.48E+04* | <LOD | <LOD | 6.20E+05 ± 8.76E+04 |
| Stationary | 0.75 | <LOD | <LOD | <LOD | <LOD | 3.85E+06 ± 3.33E+05 |
| Stationary | 0 | <LOD | <LOD | <LOD | <LOD | 5.36E+06 ± 6.77E+05 |

**Table S5** Mean attomole per cell (± SE) of cobalamins and DMB in *Thalassiosiraceae GTAN1* (n=3).

| **Growth Phase** | **Treatment** | **Ado-$B_{12}$** | **Me-$B_{12}$** | **OH-$B_{12}$** | **CN- $B_{12}$** | **Alpha-Ribazole** |
|---|---|---|---|---|---|---|
| Exponential | 100 | 0.24 ± 0.20 | 0.12 ± 0.07 | 0.28 ± 0.10 | <LOD | <LOD |
| Exponential | 10 | 0.32 ± 0.07 | 0.09 ± N/A | <LOD | <LOD | 0.60 ± 0.08 |
| Exponential | 0.75 | <LOD | <LOD | <LOD | <LOD | 1.06 ± 0.06 |
| Exponential | 0 | <LOD | <LOD | <LOD | <LOD | 1.04 ± 0.08 |
| Stationary | 100 | 0.63 ± 0.08 | 0.15 ± 0.02 | 0.47 ± 0.10 | <LOD | 0.97 ± 0.55 |
| Stationary | 10 | 0.04* | 0.09* | <LOD | <LOD | 1.03 ± 0.15 |
| Stationary | 0.75 | <LOD | <LOD | <LOD | <LOD | 6.39 ± 0.55 |
| Stationary | 0 | <LOD | <LOD | <LOD | <LOD | 8.89 ± 1.12 |

*one biological replicate above LOD

**Table S6** Shapiro-Wilk Test, Levene Test, One-Way ANOVA for the effects of cobalamin availability on DMB quotas

| Analyte | Growth Phase | Shapiro-Wilk P-value | LeveneTest P-value | Source of variation | DF | Sum Sq | Mean Sq | F value | Pr (>F) |
|---|---|---|---|---|---|---|---|---|---|
| Alpha-Ribazole | Stationary | 0.09845 | 0.3387 | Treatment | 3 | 4.502e+13 | 1.501e+13 | 15.31 | 0.001 |
| | | | | Residuals | 8 | 7.843e+12 | 9.804e+11 | | |
| | Exponential | 0.4136 | 0.7443 | Treatment | 3 | 8.094e+11 | 2.698e+11 | 43.57 | 2.66e-05 |
| | | | | Residuals | 8 | 4.954e+10 | 6.193e+9 | | |

**Table S7** Tukey Post-hoc Test for the effects of cobalamin availability on DMB quotas

| Analyte | Growth Phase | Comparison | diff | lwr | upr | p adj |
|---|---|---|---|---|---|---|
| Alpha-Ribazole | Exponential | 0.75-0 | 8.81E+03 | -1.97E+05 | 2.15E+05 | 9.99E-01 |
| | | 10-0 | -2.68E+05 | -4.74E+05 | -6.24E+04 | 1.32E-02 |
| | | 100-0 | -6.28E+05 | -8.34E+05 | -4.23E+05 | 4.67E-05 |
| | | 10-0.75 | -2.77E+05 | -4.83E+05 | -7.12E+04 | 1.10E-02 |
| | | 100-0.75 | -6.37E+05 | -8.43E+05 | -4.31E+05 | 4.21E-05 |
| | | 100-10 | -3.60E+05 | -5.66E+05 | -1.54E+05 | 2.26E-03 |
| | Stationary | 0.75-0 | -1.10E+06 | -3.69E+06 | 1.49E+06 | 5.57E-01 |
| | | 10-0 | -4.33E+06 | -6.92E+06 | -1.74E+06 | 3.03E-03 |
| | | 100-0 | -4.36E+06 | -6.95E+06 | -1.77E+06 | 2.89E-03 |
| | | 10-0.75 | -3.23E+06 | -5.82E+06 | -6.42E+05 | 1.67E-02 |
| | | 100-0.75 | -3.26E+06 | -5.85E+06 | -6.75E+05 | 1.58E-02 |
| | | 100-10 | -3.31E+04 | -2.62E+06 | 2.56E+06 | 1.00E+00 |

### Supplemental Figures

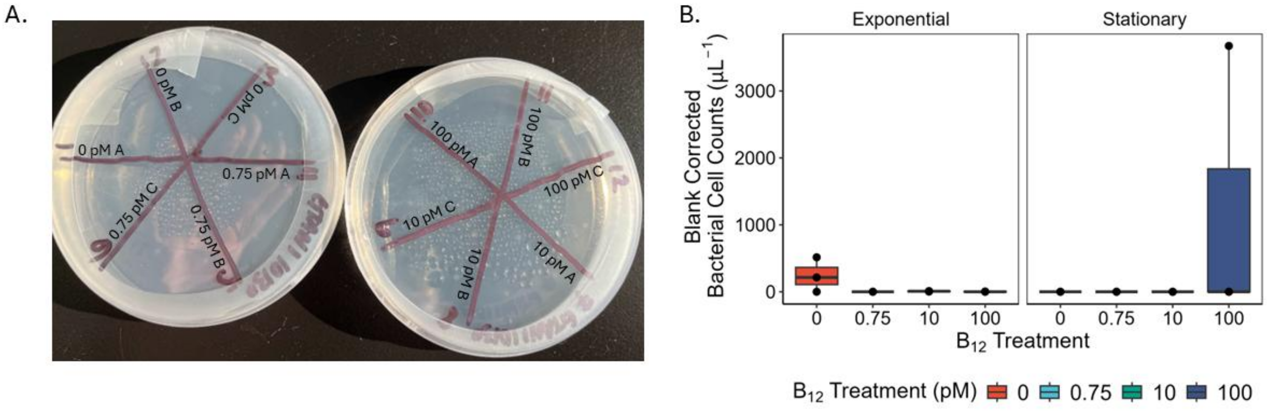

**Figure S1** Bacterial contamination assessment thourghout experiment A) Contamination check for 35 µL of culture aliquoted onto ½ Marine Broth agar plates before exponential harvest, each number represents a biological replicate . B) Blank corrected bacterial cell counts of 3 µm filtered *Thalassiosiraceae GTAN1* cultures for all $B_{12}$ treatments, samples were measured using flow cytometry. Error bars represent the standard error of biological replicates within each treatment, points represent biological replicates (n =3).

**References**


1. Guillard RRL, Ryther JH. Studies of marine planktonic diatoms: Cyclotella nana Hustedt and Detonula confervacea (Cleve) Gran. Can J Microbiol. 1962 Apr 1;8(2):229–39.

2. Heal KR, Qin W, Ribalet F, Bertagnolli AD, Coyote-Maestas W, Hmelo LR, et al. Two distinct pools of $B_{12}$ analogs reveal community interdependencies in the ocean. Proc Natl Acad Sci USA. 2017 Jan 10;114(2):364–9.

3. Bannon C, White PL, Rowland E, More KJ, Gleason A, Roberts M, et al. Seasonal patterns in b-vitamins and cobalamin co-limitation in the Northwest Atlantic [Internet]. Microbiology; 2024 [cited 2024 Nov 20]. Available from: http://biorxiv.org/lookup/doi/10.1101/2024.11.10.622835

4. Boysen AK, Heal KR, Carlson LT, Ingalls AE. Best-Matched Internal Standard Normalization in Liquid Chromatography–Mass Spectrometry Metabolomics Applied to Environmental Samples. Anal Chem. 2018 Jan 16;90(2):1363–9.

5. Paerl RW, Curtis NP, Bittner MJ, Cohn MR, Gifford SM, Bannon CC, et al. Use and detection of a vitamin B1 degradation product yields new views of the marine B1 cycle and plankton metabolite exchange. Giovannoni SJ, editor. mBio. 2023 Jun 28;e00061-23.

6. Forster R, Kromkamp J. Modelling the effects of chlorophyll fluorescence from subsurface layers on photosynthetic efficiency measurements in microphytobenthic algae. Mar Ecol Prog Ser. 2004;284:9–22.

7. Wickham H. ggplot2: Elegant Graphics for Data Analysis [Internet]. Springer-Verlag New York; 2009. Available: http://ggplot2.org
